# A method for inducing a senescence trend in mesenchymal stem cells using D-galactose^1^

**DOI:** 10.1101/2025.01.19.633537

**Authors:** Xiaoming Ji, Xiaoyu Zhou, Tingting Chen, Chunchun Duan, Wenjing Tian, Liyang Zhu, Yan Ma, Liyang Gao

**Affiliations:** Key Laboratory of Ministry of Education for Conservation and Utilization of Special Biological Re-sources in the Western, Ningxia University, Yinchuan, China; Life Science School, Ningxia University, Yinchuan, China

**Keywords:** mesenchymal stem cells, senescence, D-galactose

## Abstract

The aging of the body is accompanied by a decline in tissue function, and this decline mainly stems from the aging of tissue stem cells. Therefore, they play a crucial role in the aging of the body.This study explores the biological characteristics of senescent hUCMSCs by establishing a replicative senescence model of hUCMSCs and a rapid senescence model induced by D - galactose (D-Gal), and analyzing the proliferative capacity, cell viability, cell cycle, pluripotency, and reactive oxygen species (ROS) levels of senescent hUCMSCs.

## 1 Introduction

Bone marrow-derived mesenchymal stem cells was one of the resources of adult stem cells, and most studies focused on bone mesenchymal stem cells (BMSCs). However, the negative correlation between the survival rate and quantity of BMSCs and the age of the donor limits their applicability. Their immunomodulatory ability also decreases with age, and the invasive harm of bone marrow aspiration poses potential safety risks to patients. Therefore, human umbilical cord mesenchymal stem cells (hUCMSCs), with the advantages of easy accessibility and no ethical issues, have gradually become the seed cells intensively studied in current cell therapy.

hUCMSCs meet the minimum standards for MSCs, they highly express CD73, CD90, and CD105, while the endothelial cell surface marker CD31, hematopoietic stem cell markers CD45 and CD34, and human leukocyte antigen-DR (HLA-DR) are low-expressed or not expressed[1]. The appropriate expression of pluripotency genes Oct4, Nanog, and Sox2 in hUCMSCs endows them with the non-tumorigenic property, making hUCMSCs have great potential and research significance in tumor suppression[2–4]. On the other hand, although hUCMSCs express major histocompatibility complex class I (MHC I), they do not express major histocompatibility complex class II (MHC II), as well as co-stimulatory molecules CD40, CD80, and CD86. Moreover, they can secrete a large amount of immunosuppressive agents such as indoleamine-2,3-dioxygenase (IDO) and prostaglandin E2 (PGE2), making hUCMSCs non-immunogenic while immunomodulating [5– 7]. They have strong compatibility in an allogeneic treatment environment, and this immune characteristic has been successfully applied in human graft-versus-host disease (GVHD)[8–10]. An increasing amount of evidence indicates that hUCMSCs are receiving more and more attention in the development of modern medicine and will become a new medical approach for difficult diseases.

Aged stem cells usually show a significant increase in cell body size, a decrease in three-dimensionality, presenting a flat fried-egg shape, a decrease in cell colony-forming ability, and emerging in irregular shapes or small colony groups. This is one of the most direct predictive indicators of stem cell aging[11]. As stem cells age, small and distinct heterochromatin spots appear in their nuclei due to transcriptional inactivation, accompanied by a decrease in osteogenic differentiation ability and a greater tendency towards adipogenicity differentiation[12]. With the aging of the organism, DNA methylation decreases due to the down-regulation of DNMT1 and DNMT3B[13,14]. Therefore, aging-related DNA methylation can be used to detect cell aging[15]. In addition, the decreased expression of pluripotency-related genes OCT4, NANOG, and SOX2 during stem cell aging[16,17] and the enhanced activity of β-galactosidase[18] can all serve as markers for detecting stem cell aging.

In this study, hUCMSCs were used as experimental materials. Aged cells were obtained through continuous sub-culturing and D-galactose induction. By comparing the similarities and differences between replicative aged cells and induced-aged cells, the appropriate induction time and D-galactose treatment dose were determined. Then, the cells showing early aging phenomena after induction were analyzed.

## 2 Methodology

### 2.1 Cell culture

Cells can only be passaged when the confluence exceeds 80%. Take the cells to be passaged out of the incubator, spray the surface with alcohol, and then quickly place them in the workbench. First, discard the original culture medium and wash the cells 2-3 times with PBS to remove the residual culture medium. Add 0.5 mL of trypsin containing 0.25% EDTA. Gently shake the dish to make the trypsin spread evenly on the bottom of the dish, ensuring full contact with the cell surface. Under the microscope, after observing that the cells become round, add 0.5 mL of complete medium to terminate the digestion. Use a sterile pipette tip to gently and slowly pipette the cell solution to blow-wash the bottom of the culture dish, trying to make all the cells detach. Transfer the digested cell solution to a 15 mL centrifuge tube. Wash the culture dish 1-2 times with PBS and collect the cell suspension into the same centrifuge tube. Centrifuge to collect the cells under the conditions of 1000 rpm for 5 minutes.

After centrifugation, discard the supernatant and add an appropriate amount of fresh serum-free complete cell culture medium to resuspend the cells. Pipette 10 μL of the cell suspension onto a cell counting chamber and count the cells under a microscope or using an automatic cell counter. According to the requirements of subsequent experiments, select an appropriate number of cells to be seeded into a culture dish containing fresh serum-free complete medium. Place the dish in a constant temperature incubator at 37 °C with 5% CO_2_ for cultivation.

### 2.2 SA-β-gal staining

Inoculate hUCMSCs at an appropriate density into a 35 mm dish and place it in a constant-temperature incubator at 37 °C with 5% CO_2_. When the detection time arrives, aspirate the old culture medium and wash it once with PBS. Add 1 mL of β-galactosidase staining fixative and fix at room temperature for 15 minutes. Immediately after fixation, aspirate the fixative and wash it 3 times with PBS, 3 minutes each time. Meanwhile, prepare the staining working solution in proportion (Staining Solution A: Staining Solution B: Staining Solution C: X-Gal Solution = 10μL: 10μL: 930μL: 50μL). After aspirating the PBS, add 1 mL of the staining working solution. Seal the mouth of the culture dish with parafilm to prevent the evaporation of the staining solution, and place the culture dish in a 37 °C incubator for overnight incubation. The next day, observe, take photos, and count under an optical microscope.

### 2.3 Detection of Reactive Oxygen Species (ROS)

Seed cells in a 6-well plate, with 3 replicate wells per group. Culture the cells until the confluence reaches 80-90%. Aspirate the culture medium and wash the cells 2-3 times with PBS. Add 1 mL of fresh complete medium to each well and add 2μL of CellROX Oxidative Stress Reagents staining solution. Incubate at 37 °C in the dark for 20 minutes. Discard the staining solution, digest the cells as in the cell passage method, and centrifuge to collect the cells. Soak and wash the cells 2-3 times with PBS solution, then add 500 μL of PBS to resuspend the cells for machine detection.

### 2.4 Detect specific marker of MSCs by flow cytometry

When the cells are passaged to the corresponding generation, centrifuge to obtain hUCMSCs cell pellets and discard the supernatant. Add 1 mL of cold PBA (PBS containing 5% FBS) to the centrifuge tube to resuspend the cells. After centrifugation, discard the supernatant. Add 200μL of fluorescein-labeled antibodies (CD90, CD73, and CD105) diluted with PBA, mix well, and incubate at 4 °C for 30-60 minutes. After the incubation, centrifuge at 1000 rpm for 5 minutes and discard the supernatant. Add 1 mL of cold PBS to the centrifuge tube to resuspend the cells, and centrifuge and wash the cells twice to remove the unbound excess antibodies. Add 500μL of cold PBS, mix well, and perform machine detection.

### 2.5 Immunofluorescence

Inoculate hUCMSCs with good growth status in a 24-well plate. Cultivate the cells with a medium containing D-Gal until the required time for the experiment. Aspirate the old culture medium in the well plate, and wash the cell climbing sheets with PBS to remove the residual culture medium. Add an appropriate amount of 4% paraformaldehyde to the well plate. Fix the cells at room temperature for 40 minutes. Then remove the fixative, and rinse the cell climbing sheets with PBS for 3 times, 5 minutes each time. Add 0.3% Triton X-100 prepared with PBS, and permeabilize the cells at room temperature for 15 minutes. Remove the permeabilization solution, and wash the cell climbing sheets with PBS for 3 times, 5 minutes each time. Add an appropriate amount of primary antibody dilution to the well plate, and block at room temperature for 30 minutes. Aspirate the blocking solution, and add the diluted primary antibodies OCT4 (1:200) and NANOG (1:800). Incubate overnight at 4 °C. Recover the primary antibody dilution, wash the cell climbing sheets with PBS, add the diluted fluorescent secondary antibody onto the climbing sheets, and incubate at room temperature in the dark for 1 hour. Wash the cell climbing sheets with PBS for 3 times, 5 minutes each time. Drop an appropriate amount of DAPI on a glass slide. After taking out the cell climbing sheet, cover it on the glass slide. Evenly smear the mounting medium along the edge of the cell climbing sheet for mounting. Place it in a dark box to dry, and then collect images using a fluorescence microscope.

### 2.6 Data Analysis

All data in this paper are obtained from more than three repeated experiments. For data analysis, IBM SPSS Statistics 22 software and GraphPad Prism 7 application software are selected, and statistical analysis is carried out using One-Way ANOVA (One-way Analysis of Variance) and T test. Data are presented as mean ± standard deviation (X ± SD). Among them, *, **, and *** all indicate significant differences, representing 0.01 < *P* < 0.05, 0.001 < *P* < 0.01, and *P* < 0.001, respectively.

## 3 Results

### 3.1 Morphology of Replicative Senescent hUCMSCs and D-galactose-induced Senescent hUCMSCs

In this experiment, hUCMSCs were selected as the research object. Under the microscope, normal hUCMSCs appear as adherent cells with an elongated spindle-shaped morphology similar to fibroblasts. When the cells grow to an appropriate density, they will show a spiral shape (Figure 1A). When continuously passaged to the 25th generation (P25) to induce replicative senescence, the cell morphology becomes flat, like a fried egg, the cell body enlarges, and the cell colony-forming ability decreases (Figure 1B).

**Fig 1.**
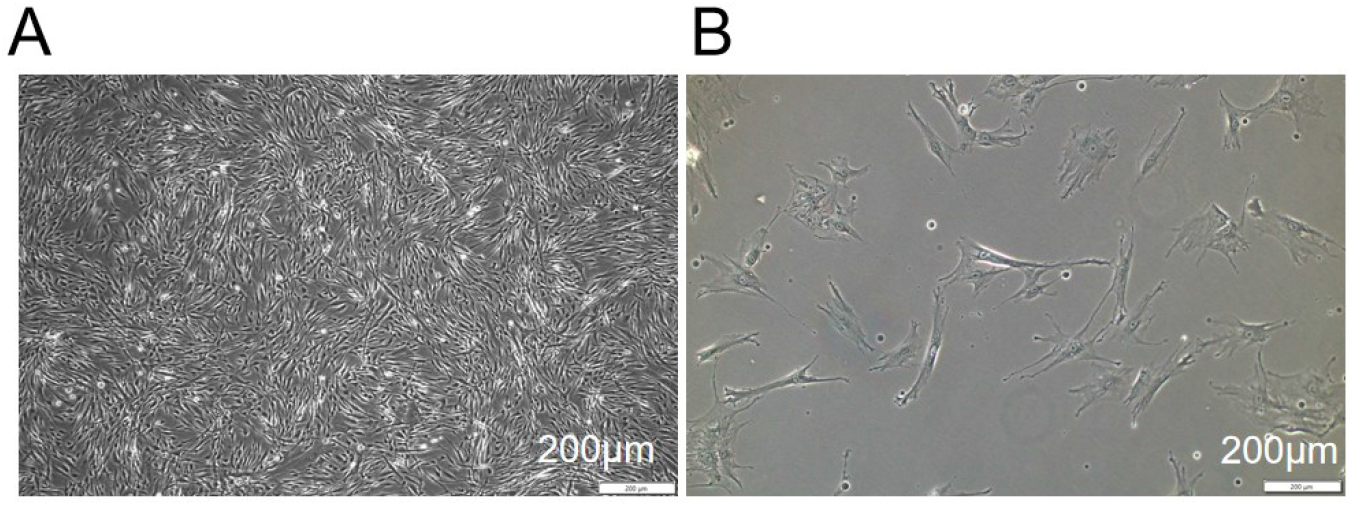
Morphological observation of normal and replicative senescent hUCMSCs A: Morphology and growth status of normal hUCMSCs; B: Morphology and growth status of senescent hUCMSCs at passage 25.

### 3.2 Senescence – related markers: Replicative Senescent hUCMSCs and D-galactose-induced Senescent hUCMSCs

#### 3.2.1 Intracellular expression of SA-β-gal in two groups

SA-β-gal detection is a powerful indicator commonly used to detect senescence. Senescent hUCMSCs are stained blue-green. In this experiment, hUCMSCs passaged to P25 were stained with SA-β-gal to detect the senescence status. The results showed that compared with normal cells, replicative senescent hUCMSCs had a higher positive staining rate (Figure 2).

**Fig 2.**
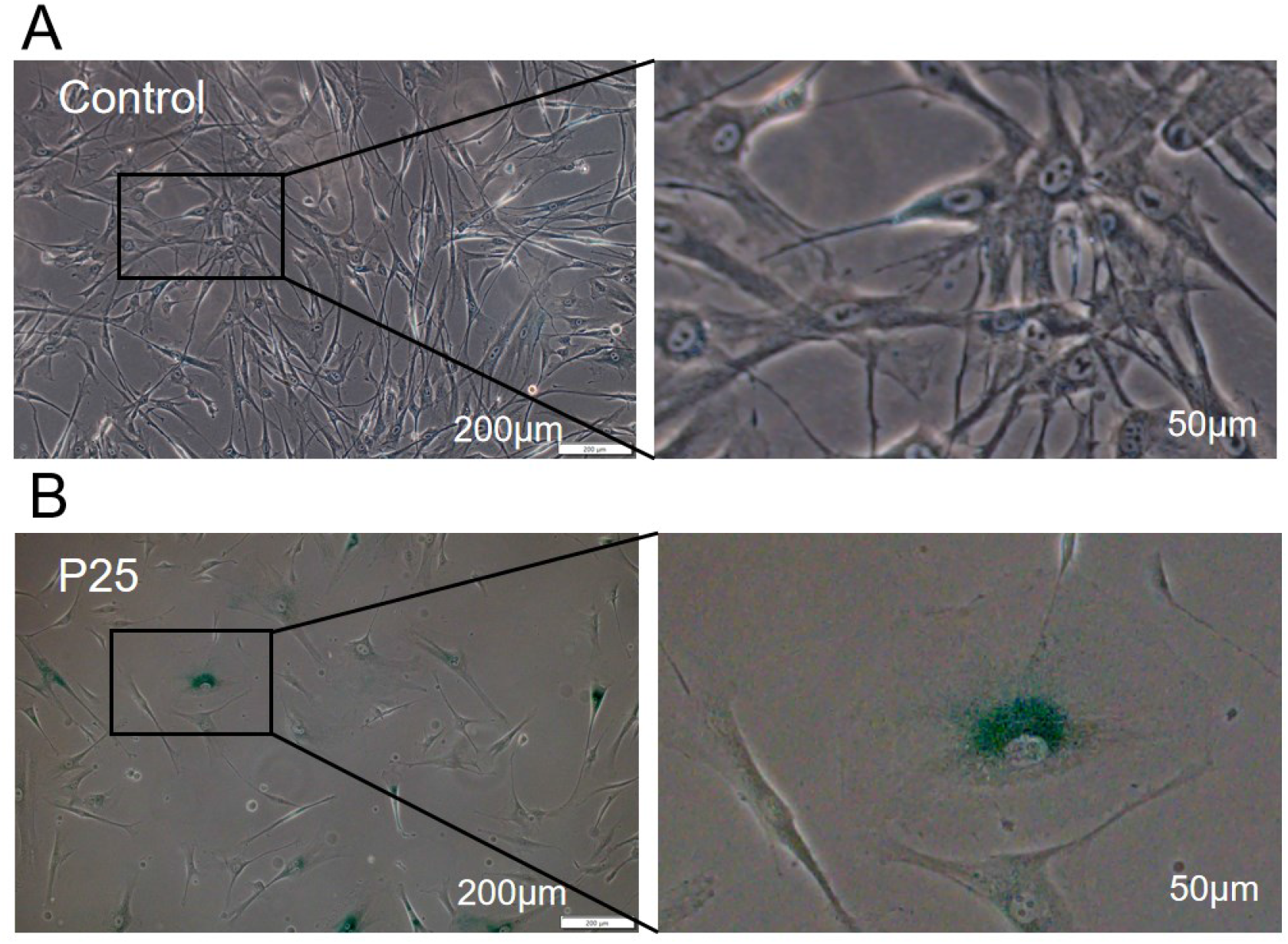
hUCMSCs were continuously passaged to P25 SA-β-gal staining

After treating hUCMSCs with 2% D-Gal for 2 days (D2) and 4 days (D4), the positive reaction of SA-β-gal staining was detected. The results showed that 2% D-Gal could effectively induce the senescence of hUCMSCs, and the positive staining rate of SA-β-gal increased with the prolongation of treatment time (Figure 3).

**Fig 3.**
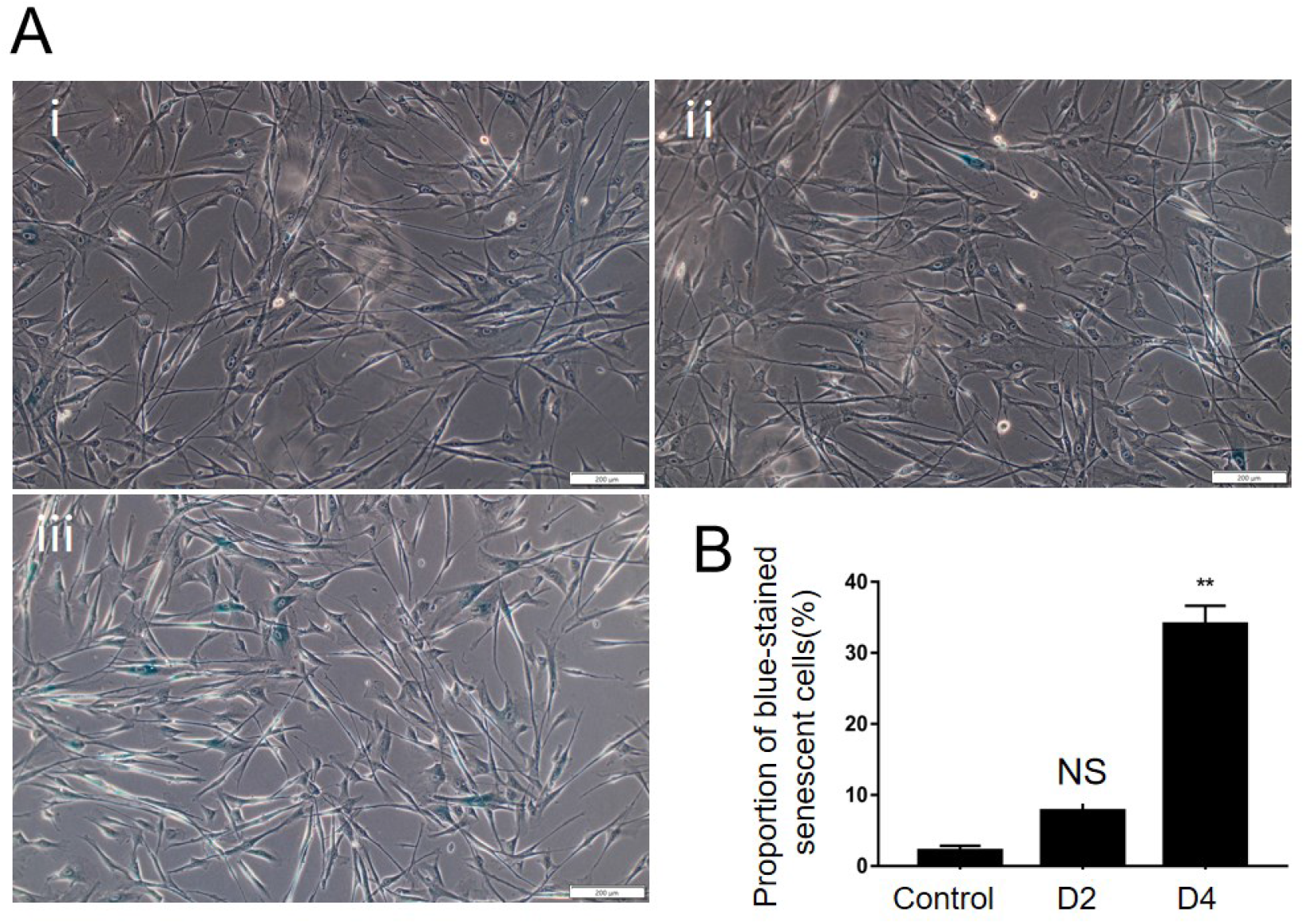
hUCMSCs were treated with 2%D-Gal and stained with SA-β-gal

#### 3.2.2 ROS level

DNA damage is generally regarded as the main cause of cell senescence and apoptosis, and endogenous ROS is the primary driving force leading to DNA damage. In view of this, in this study, an oxidative stress fluorescent probe was used to detect the ROS levels in normal hUCMSCs and replicative senescent hUCMSCs that had been continuously passaged to P25 by flow cytometry. The results showed that the ROS level in replicative senescent hUCMSCs increased significantly (Figure 4), and ROS accumulated in hUCMSCs as the cells were passaged.

**Fig 4.**
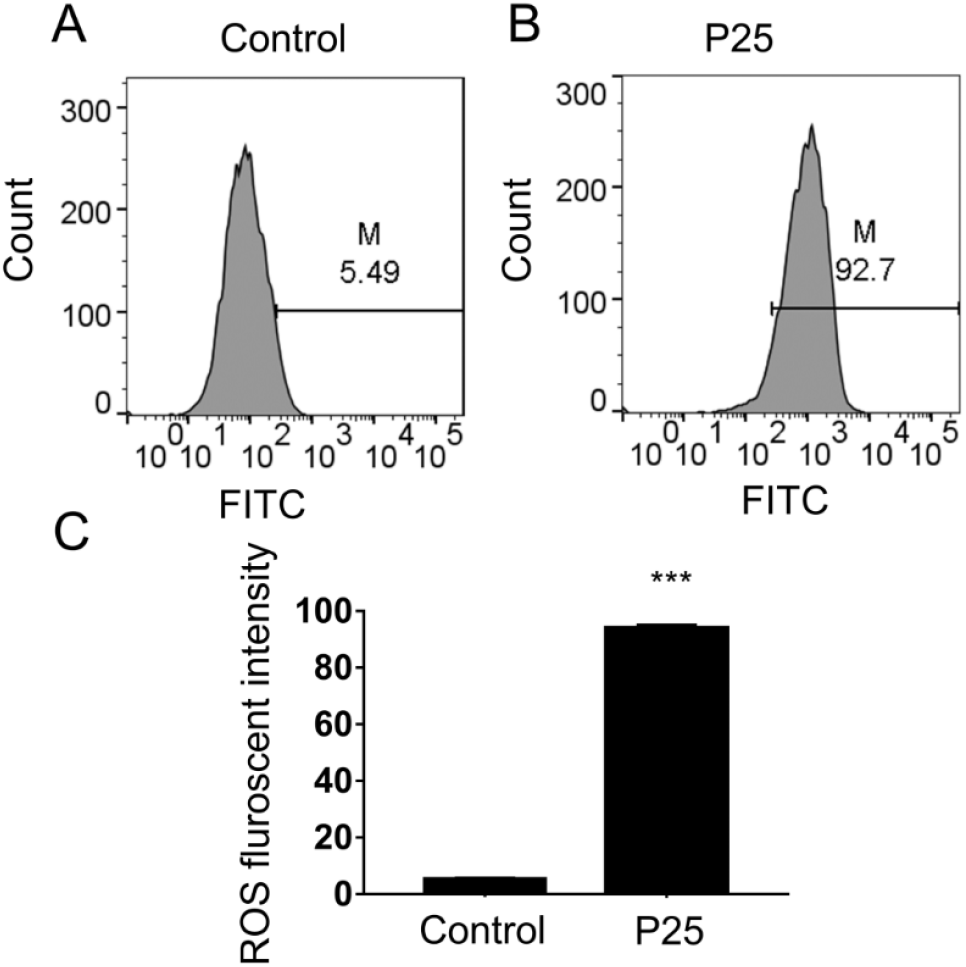
ROS detection of replicative senescent hUCMSCs A: Detection of ROS level in normal hUCMSCs; B: Detection of ROS level in senescent cells at passage 25; C: Comparison of ROS levels between senescent cells at passage 25 and cells in the normal group.*:0.01<*P*<0.05,**:0.001<*P*<0.01,***:*P*<0.001

To determine whether the senescence of hUCMSCs induced by D-Gal leads to an increase in the ROS level, in this experiment, the ROS levels in hUCMSCs treated with 2% D-Gal for 2 days (D2) and 4 days (D4) were detected. The results showed that the intracellular ROS level increased with the increase of treatment time, and the increase was significant after 4 days of treatment (Figure 5). Moreover, this result was consistent with that of the replicative senescence model.

**Fig 5.**
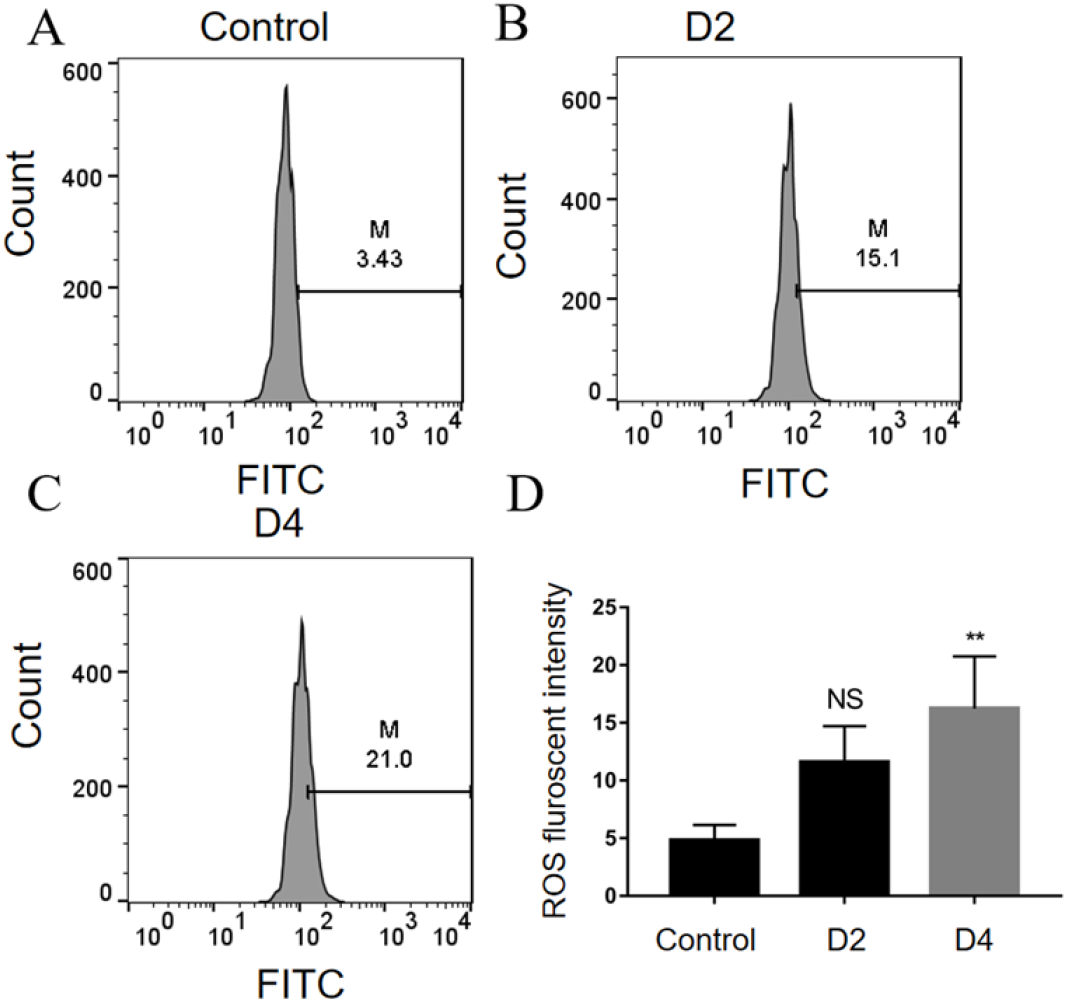
Effects of 2%D-Gal treatment on ROS in hUCMSCs

#### 3.2.3 Mesenchymal stem cell-specific surface markers

Flow cytometry was used to detect the surface molecular markers of normal hUCMSCs and hUCMSCs at passage 25 (P25). It was found that both groups expressed CD90, CD73, and CD105. Although replicative senescent hUCMSCs possessed typical mesenchymal stem cell surface molecular markers, their expression levels were lower than those of the normal group (Figure 6). The results indicated that replicative senescent hUCMSCs obtained through continuous passage gradually lost their pluripotency.

**Fig 6.**
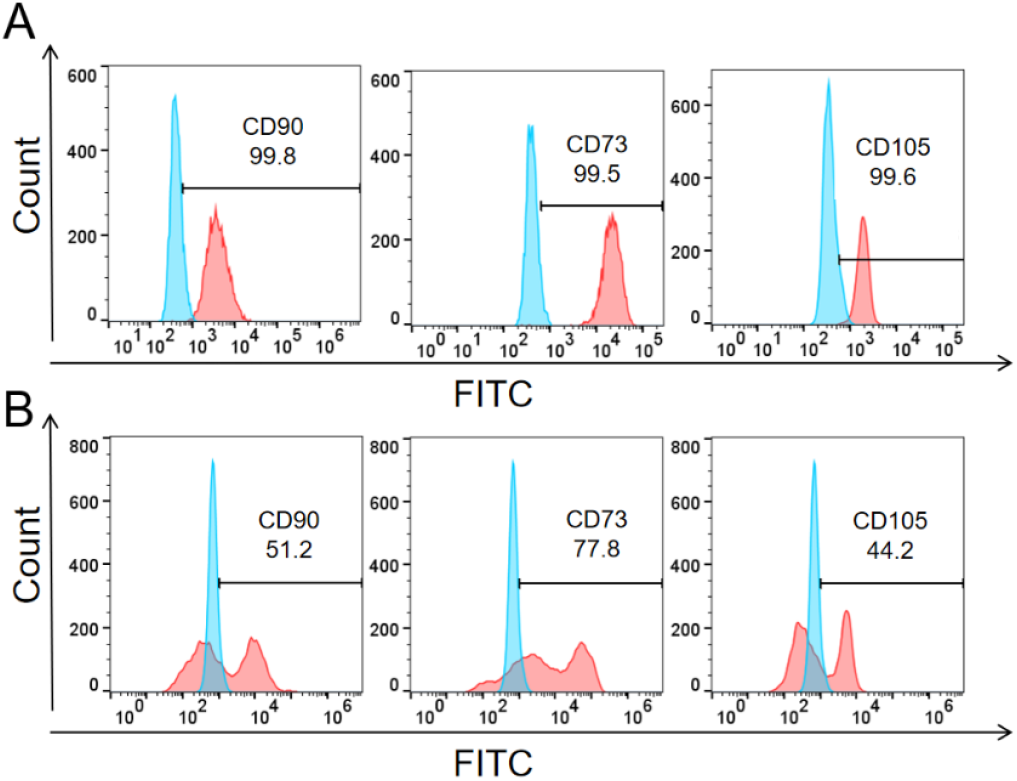
Detection of Surface Markers of Normal and replicative Senescent hUCMSCs (Flow Cytometry). A: Flow cytometry detection of the expression of pluripotency - related proteins in normal hUCMSCs. B: Flow cytometry detection of the expression of pluripotency - related proteins in replicatively senescent hUCMSCs at passage 25.

To assess the effect of 2% D-Gal on inducing cell senescence, flow cytometry was employed to detect the expression of the marker proteins CD90, CD73, and CD105 in hUCMSCs treated with 2% D-Gal for 2 days (D2), 4 days (D4), and 6 days (D6). The results showed that all three marker proteins were positive, with the expression of CD90 and CD73 exceeding 99%. The expression of CD105 decreased with the treatment time, and the positive rate was only 88.6% after 6 days of treatment (Figure 7).

**Fig 7.**
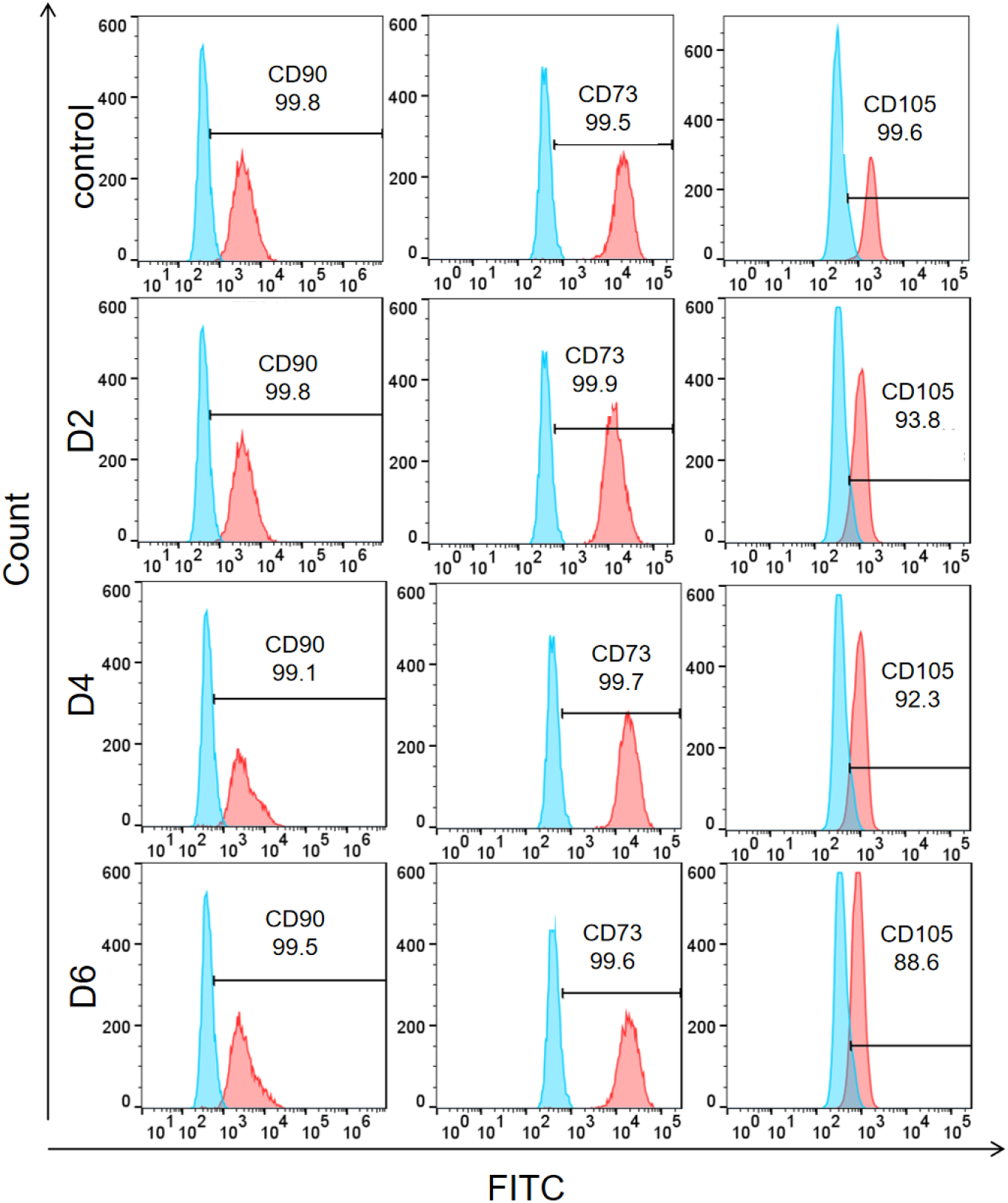
Flow cytometric detection of marker proteins in hUCMSCs treated with 2%D-Gal A:i: SA - β - gal staining of hUCMSCs without D - Gal treatment; ii: SA - β - gal staining of hUCMSCs after 2 - day (D2) treatment with D - Gal. Senescent cells appear blue - green; iii: SA - β - gal staining of hUCMSCs after 4 - day (D4) treatment with D - Gal. Senescent cells appear blue - green. B: Comparison of the percentage differences of senescent cells in hUCMSCs after different treatment times with D – Gal. *:0.01<*P*<0.05,**:0.001<*P*<0.01,***:*P*<0.001

#### 3.3 Effect of 2% D-Gal Induction on the Proliferation of hUCMSCs

To investigate whether 2% D-Gal affects the cellular functions of hUCMSCs, this experiment used the CCK-8 method to detect the effect of 2% D-Gal on their cell proliferation ability (Figure 8). The results showed that the cell proliferation ability of hUCMSCs significantly decreased after 4 days of treatment with 2% D-Gal. The CellTiter-LumiTM luminescence method was used to detect the effect of 2% D-Gal on the cell viability of hUCMSCs (Figure 8). The results indicated that 2% D-Gal could significantly reduce the cell viability of hUCMSCs. The above results suggest that 2% D-Gal can effectively decrease the cell proliferation ability and cell viability of hUCMSCs. To further determine the impact of D-Gal on the cell proliferation of hUCMSCs, this experiment detected the cell cycle of hUCMSCs after treatment with 2% D-Gal by flow cytometry (Figure 8). The results showed that hUCMSCs treated with 2% D-Gal for 4 days exhibited a G0/G1-phase cell-cycle arrest, mainly manifested as an increase in the percentage of cells in the G1 phase and a decrease in the percentage of cells in the G2 phase.

**Fig 8.**
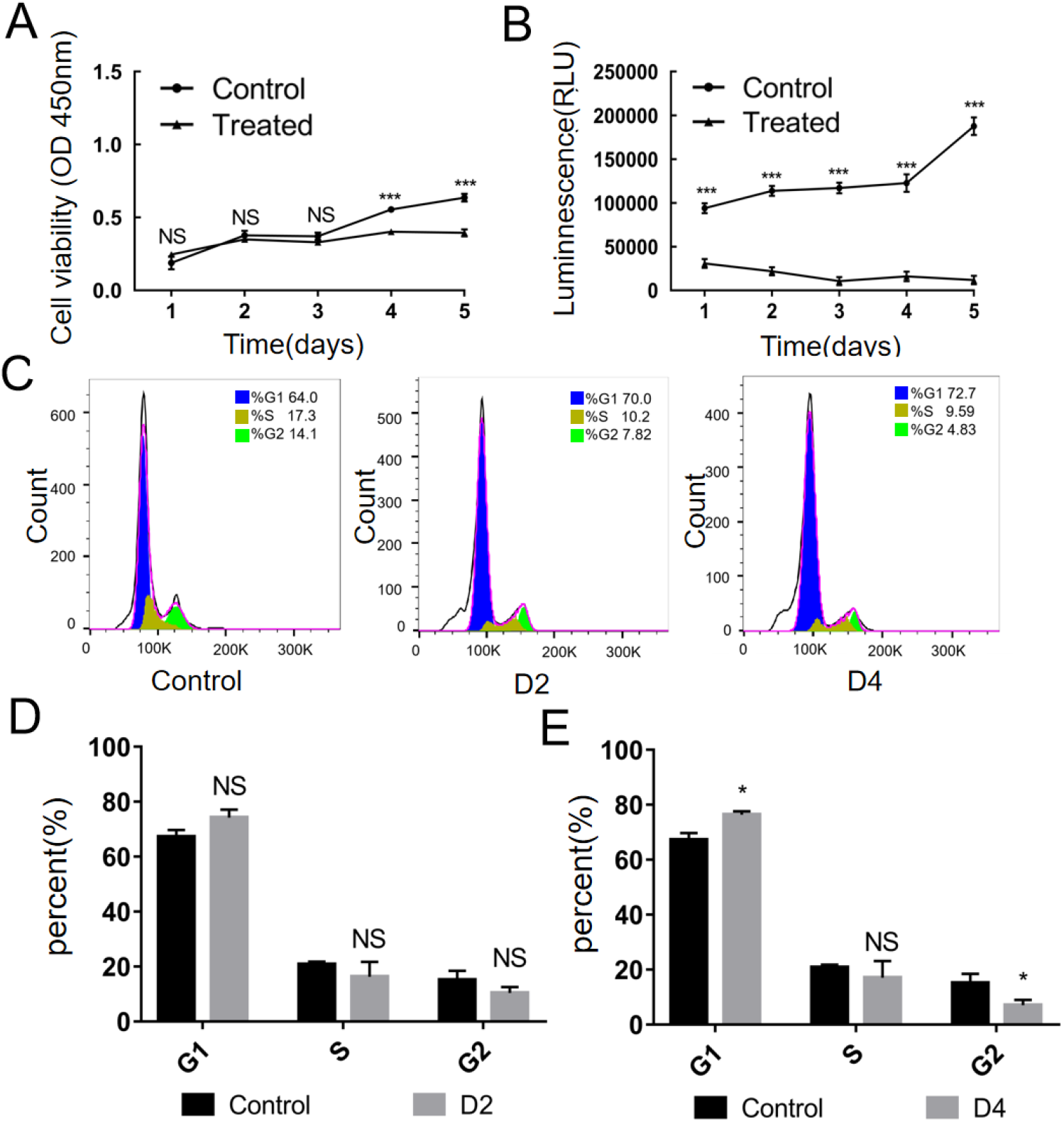
Effect of 2%D-Gal on hUCMSCs cell function A: Detection of the effect of 2% D - Gal treatment on the cell proliferation ability of hUCMSCs by the CCK - 8 method. B: Detection of the effect of 2% D - Gal on the cell viability of hUCMSCs by the CellTiter – LumiTM luminescence method. C - E: Detection of the effect of 2% D - Gal on the cell cycle of hUCMSCs by flow cytometry. *:0.01<*P*<0.05,**:0.001<*P*<0.01,***:*P*<0.001

#### 3.3 Effect of 2% D-Gal on the Pluripotency of hUCMSCs

To explore whether 2% D-Gal could lead to a decline in the pluripotency of hUCMSCs, in this experiment, the expression of pluripotency-related genes C-KIT, NANOG, and OCT4 in hUCMSCs without D-galactose treatment, as well as those treated with 2% D-Gal for 2 days (Figure 9) and 4 days (Figure 6), was detected by chemico-immunofluorescence. The results showed that hUCMSCs still expressed the pluripotency-related proteins C-KIT, NANOG, and OCT4 after being treated with 2% D-Gal for 2 days and 4 days. Western blot was used to detect whether the expression of NANOG and OCT4 was affected after hUCMSCs were treated with 2% D-Gal for 2 days and 4 days. The results showed that the expression of OCT4 was significantly upregulated with the increase of D-Gal treatment time, while the expression of NANOG was significantly downregulated after 4 days of D-Gal treatment (Figure 9). This indicates that D-Gal has a certain impact on the pluripotency of hUCMSCs, but this impact is not simply to make them lose pluripotency. The upregulation of OCT4 expression is considered as a regulatory way for hUCMSCs to resist senescence, which is a phenomenon of early senescence.

**Fig 9.**
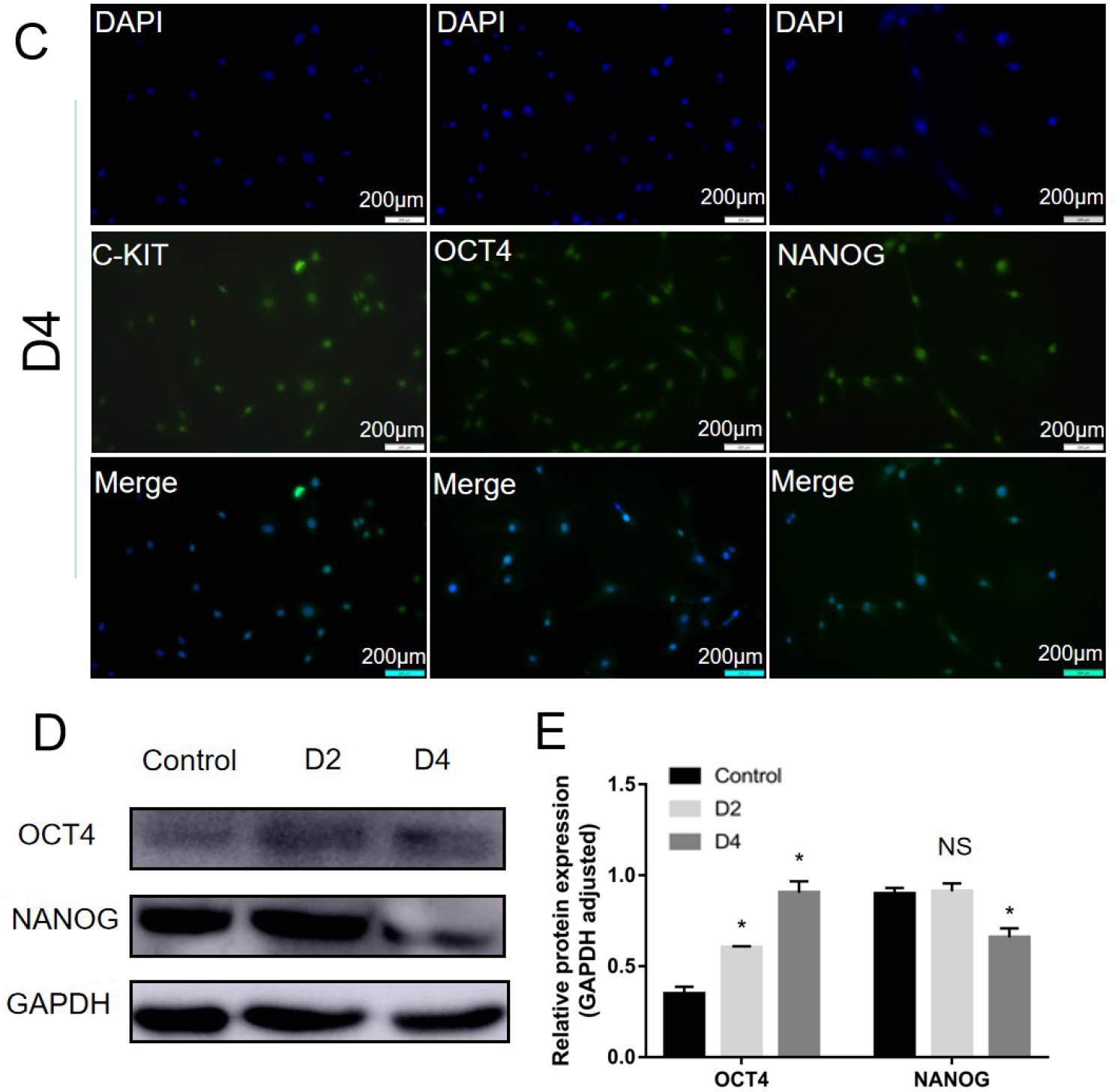
Effects of 2%D-Gal on genes related to pluripotency in hUCMSCs 2 and 4 days after treatment. A: Detection of the expression of pluripotency - related genes in hUCMSCs without; D - Gal treatment by immunofluorescence.; B: Detection of the expression of pluripotency - related genes in hUCMSCs treated with 2% D - Gal for 2 days by immunofluorescence.; C: Detection of the expression of pluripotency - related genes in hUCMSCs treated with 2% D - Gal for 4 days by immunofluorescence. D - E: Detection of the expression of pluripotency - related proteins in hUCMSCs treated with 2% D - Gal for 2 days and 4 days by Western Blot.*:0.01<*P*<0.05,**:0.001<*P*<0.01,***:*P*<0.001

## 4. Discussion

Cell senescence can be classified into replicative senescence mediated by an increase in the number of cell passages and premature senescence induced by stress from exogenous substances. During the study of cell senescence, the establishment of a replicative senescence model is time-consuming and costly. Therefore, the stress-induced accelerated senescence model is more favored due to its advantages such as simple application, short research time, and low cost. Common accelerated senescence models include D-Gal[19], H_2_O_2_[20,21], radiation[22–24], etc. Among these senescence models, the D-Gal-induced senescence model is a preferred inducer because of its convenience and low side-effects.

D-Gal, an aldose, naturally exists in the body and many foods. For healthy adults, the maximum recommended daily dose of galactose is 50g. High levels of galactose can be converted into aldose and hydroperoxide under the catalysis of galactose oxidase, subsequently generating reactive oxygen species (ROS), which then lead to inflammation, oxidative stress, mitochondrial dysfunction, and apoptosis[25–28]. Research shows that D-Gal induces an increase in senescence markers, such as sorbitol dehydrogenase (SDH), advanced glycation end-products (AGE), aldose reductase (AR), receptor for advanced glycation end-products (RAGE), telomerase activity, telomere shortening, β-site amyloid-precursor protein-cleaving enzyme 1 (BACE-1), senescence-associated genes (P16, P21, P53, P19Arf, P21Cip1/Waf1), β-amyloid protein, and senescence-associated SA-β-gal staining[29]. Several in vitro studies have utilized the D-Gal-induced accelerated senescence model on different cell types, such as primary neural stem cells (NSCs), astrocytes, stromal cells, human umbilical vein endothelial cells, and renal proximal tubular epithelial cells. In the NSC D-Gal senescence model, the levels of malondialdehyde (MDA) and ROS increased significantly, and the decrease in superoxide dismutase (SOD) levels and total antioxidant capacity indicated the occurrence of an oxidative stress response, accompanied by an increase in the levels of inflammatory factors such as TNF-α, IL-6, IL-1β, etc., and the significant expression of P53 and P21. This proves that D-Gal may induce senescence through the inflammatory pathway and the P53-P21 pathway[30,31]. In the astrocyte aging model established by D-Gal, similarities were found between D-Gal-induced senescence and naturally occurring senescence, but age-related mitochondrial energy metabolism, glycolytic activity, and the glutamate-glutamine cycle were not completely consistent in the two types of senescence. D-Gal-induced astrocytes showed a decrease in basal glycolytic activity, while the opposite was true in replicative senescence. The iron-sulfur cluster assembly enzyme (ISCU) was upregulated in replicative senescence but downregulated in the D-Gal-induced model. The protein levels of phosphoinositide 3-kinase (PI3K), phosphorylated AKT (p-AKT), tyrosine-kinase B (AKT), and phosphorylated glycogen synthase kinase 3β (p-GSK3β) decreased in replicative senescent cells, but only the reduction of PI3K and p-AKT was observed in the D-Gal senescence model. These differences are speculated to be due to the different degrees of aging of D-Gal-induced and naturally occurring senescent astrocytes[30]. In the mouse testicular cell senescence model induced by different concentrations of D-Gal, D-Gal significantly decreased cell viability in a concentration-dependent manner and showed cell-cycle arrest in the G0/G1 phase[32].

## 5. Conclusion

In this study, D-Gal was used to induce hUCMSCs to construct a rapid senescence model. The induction concentration of D-Gal was determined by the cell proliferation performance of cells treated with D-Gal at concentrations of 2% (2g/100mL) and 4% (4g/100mL). The induction time was determined by SA-β-gal staining. By comparing the senescent cells obtained from the constructed hUCMSCs rapid senescence model in terms of pluripotency, cell cycle, ROS, etc., it was found that their biological characteristics were similar to those of replicative senescent cells. However, replicative senescent cells lost more pluripotency and accumulated more ROS. This is considered to be due to the fact that the degree of senescence of hUCMSCs induced by D-Gal is weaker than that of replicative senescence, belonging to early senescence. Compared with replicative senescence, this model has the advantages of short time, low cost, and simple operation, and can replace the replicative senescence model in the study of hUCMSCs senescence. However, whether their internal mechanisms are consistent remains to be verified.

## Acknowledgments

This work was supported by Ningxia Natural Science Foundation (2023AAC03118), National Natural Science Foundation of China (31960189), and Ningxia Key R&D Plan Project (2018BFH03018).

## Author Contributions

Conceptualization, Liyang Gao; methodology, Xiaoming Ji, Xiaoyu Zhou; validation, Liyang Gao, TingTing Chen.; formal analysis, Liyang Gao, Wenjing Tian; data curation, Xiaoming Ji, Xiaoyu Zhou; writing—original draft preparation, Xiaoyu Zhou, Xiaoming Ji; writing—review and editing, Liyang Gao, Chunchun Duan, Liyang Zhu, Yan Ma; funding acquisition, Liyang Gao. All authors have read and agreed to the published version of the manuscript.

## Data Availability Statement

The original contributions presented in the study are included in the article/Supplementary Material.

## Reference

1. Venugopal P, Balasubramanian S, Majumdar A Sen, Ta M. Isolation, characterization, and gene expression analysis of Wharton’s jelly-derived mesenchymal stem cells under xeno-free culture conditions. Stem Cells Cloning. 2011;4: 39–50. doi:10.2147/SCCAA.S17548

2. Meng M-Y, Li L, Wang W-J, Liu F-F, Song J, Yang S-L, et al. Assessment of tumor promoting effects of amniotic and umbilical cord mesenchymal stem cells in vitro and in vivo. J Cancer Res Clin Oncol. 2019;145: 1133–1146. doi:10.1007/s00432-019-02859-6

3. Piccinato CA, Sertie AL, Torres N, Ferretti M, Antonioli E. High OCT4 and Low p16(INK4A) Expressions Determine In Vitro Lifespan of Mesenchymal Stem Cells. Stem Cells Int. 2015;2015: 369828. doi:10.1155/2015/369828

4. Slama Y, Ah-Pine F, Khettab M, Arcambal A, Begue M, Dutheil F, et al. The Dual Role of Mesenchymal Stem Cells in Cancer Pathophysiology: Pro-Tumorigenic Effects versus Therapeutic Potential. Int J Mol Sci. 2023;24. doi:10.3390/ijms241713511

5. Stefańska K, Ożegowska K, Hutchings G, Popis M, Moncrieff L, Dompe C, et al. Human Wharton’s Jelly-Cellular Specificity, Stemness Potency, Animal Models, and Current Application in Human Clinical Trials. J Clin Med. 2020;9. doi:10.3390/jcm9041102

6. Zhang Z, Huang S, Wu S, Qi J, Li W, Liu S, et al. Clearance of apoptotic cells by mesenchymal stem cells contributes to immunosuppression via PGE2. EBioMedicine. 2019;45: 341–350. doi:10.1016/j.ebiom.2019.06.016

7. Mishra VK, Shih H-H, Parveen F, Lenzen D, Ito E, Chan T-F, et al. Identifying the Therapeutic Significance of Mesenchymal Stem Cells. Cells. 2020;9. doi:10.3390/cells9051145

8. Li Y, Hao J, Hu Z, Yang Y-G, Zhou Q, Sun L, et al. Current status of clinical trials assessing mesenchymal stem cell therapy for graft versus host disease: a systematic review. Stem Cell Res Ther. 2022;13: 93. doi:10.1186/s13287-022-02751-0

9. Keshavarz Shahbaz S, Mansourabadi AH, Jafari D. Genetically engineered mesenchymal stromal cells as a new trend for treatment of severe acute graft-versus-host disease. Clin Exp Immunol. 2022;208: 12–24. doi:10.1093/cei/uxac016

10. Markov A, Thangavelu L, Aravindhan S, Zekiy AO, Jarahian M, Chartrand MS, et al. Mesenchymal stem/stromal cells as a valuable source for the treatment of immune-mediated disorders. Stem Cell Res Ther. 2021;12: 192. doi:10.1186/s13287-021-02265-1

11. Hong X, Wang L, Zhang K, Liu J, Liu J-P. Molecular Mechanisms of Alveolar Epithelial Stem Cell Senescence and Senescence-Associated Differentiation Disorders in Pulmonary Fibrosis. Cells. 2022;11. doi:10.3390/cells11050877

12. Abuna RPF, Stringhetta-Garcia CT, Fiori LP, Dornelles RCM, Rosa AL, Beloti MM. Aging impairs osteoblast differentiation of mesenchymal stem cells grown on titanium by favoring adipogenesis. J Appl Oral Sci. 2016;24: 376–382. doi:10.1590/1678-775720160037

13. Cakouros D, Gronthos S. Epigenetic Regulation of Bone Marrow Stem Cell Aging: Revealing Epigenetic Signatures associated with Hematopoietic and Mesenchymal Stem Cell Aging. Aging Dis. 2019;10: 174–189. doi:10.14336/AD.2017.1213

14. Singh P, Kacena MA, Orschell CM, Pelus LM. Aging-Related Reduced Expression of CXCR4 on Bone Marrow Mesenchymal Stromal Cells Contributes to Hematopoietic Stem and Progenitor Cell Defects. Stem cell Rev reports. 2020;16: 684–692. doi:10.1007/s12015-020-09974-9

15. Guerrero EN, Vega S, Fu C, De León R, Beltran D, Solis MA. Increased proliferation and differentiation capacity of placenta-derived mesenchymal stem cells from women of median maternal age correlates with telomere shortening. Aging (Albany NY). 2021;13: 24542–24559. doi:10.18632/aging.203724

16. Barbet R, Peiffer I, Hatzfeld A, Charbord P, Hatzfeld JA. Comparison of Gene Expression in Human Embryonic Stem Cells, hESC-Derived Mesenchymal Stem Cells and Human Mesenchymal Stem Cells. Stem Cells Int. 2011;2011: 368192. doi:10.4061/2011/368192

17. Chen H, Liu X, Zhu W, Chen H, Hu X, Jiang Z, et al. SIRT1 ameliorates age-related senescence of mesenchymal stem cells via modulating telomere shelterin. Front Aging Neurosci. 2014;6: 103. doi:10.3389/fnagi.2014.00103

18. de Mera-Rodríguez JA, Álvarez-Hernán G, Gañán Y, Martín-Partido G, Rodríguez-León J, Francisco-Morcillo J. Is Senescence-Associated β-Galactosidase a Reliable in vivo Marker of Cellular Senescence During Embryonic Development? Front cell Dev Biol. 2021;9: 623175. doi:10.3389/fcell.2021.623175

19. Wang J, Liu L, Ding Z, Luo Q, Ju Y, Song G. Exogenous NAD(+) Postpones the D-Gal-Induced Senescence of Bone Marrow-Derived Mesenchymal Stem Cells via Sirt1 Signaling. Antioxidants (Basel, Switzerland). 2021;10. doi:10.3390/antiox10020254

20. Pieńkowska N, Bartosz G, Pichla M, Grzesik-Pietrasiewicz M, Gruchala M, Sadowska-Bartosz Effect of antioxidants on the H(2)O(2)-induced premature senescence of human fibroblasts. Aging (Albany NY). 2020;12: 1910–1927. doi:10.18632/aging.102730

21. Yan J, Wang J, Huang H, Huang Y, Mi T, Zhang C, et al. Fibroblast growth factor 21 delayed endothelial replicative senescence and protected cells from H(2)O(2)-induced premature senescence through SIRT1. Am J Transl Res. 2017;9: 4492–4501.

22. Wang Y, Xu L, Wang J, Bai J, Zhai J, Zhu G. Radiation induces primary osteocyte senescence phenotype and affects osteoclastogenesis in vitro. Int J Mol Med. 2021;47. doi:10.3892/ijmm.2021.4909

23. Day RM, Snow AL, Panganiban RAM. Radiation-induced accelerated senescence: a fate worse than death? Cell cycle (Georgetown, Tex.). United States; 2014. pp. 2011–2012. doi:10.4161/cc.29457

24. Kim JH, Brown SL, Gordon MN. Radiation-induced senescence: therapeutic opportunities. Radiat Oncol. 2023;18: 10. doi:10.1186/s13014-022-02184-2

25. Yang M, Teng S, Ma C, Yu Y, Wang P, Yi C. Ascorbic acid inhibits senescence in mesenchymal stem cells through ROS and AKT/mTOR signaling. Cytotechnology. 2018;70: 1301–1313. doi:10.1007/s10616-018-0220-x

26. Al-Azab M, Wang B, Elkhider A, Walana W, Li W, Yuan B, et al. Indian Hedgehog regulates senescence in bone marrow-derived mesenchymal stem cell through modulation of ROS/mTOR/4EBP1, p70S6K1/2 pathway. Aging (Albany NY). 2020;12: 5693–5715. doi:10.18632/aging.102958

27. Fan C, Ji Q, Zhang C, Xu S, Sun H, Li Z. TGF-β induces periodontal ligament stem cell senescence through increase of ROS production. Mol Med Rep. 2019;20: 3123–3130. doi:10.3892/mmr.2019.10580

28. Vigneron A, Vousden KH. p53, ROS and senescence in the control of aging. Aging (Albany NY). 2010;2: 471–474. doi:10.18632/aging.100189

29. Shwe T, Pratchayasakul W, Chattipakorn N, Chattipakorn SC. Role of D-galactose-induced brain aging and its potential used for therapeutic interventions. Exp Gerontol. 2018;101: 13– 36. doi:10.1016/j.exger.2017.10.029

30. Sha J-Y, Li J-H, Zhou Y-D, Yang J-Y, Liu W, Jiang S, et al. The p53/p21/p16 and PI3K/Akt signaling pathways are involved in the ameliorative effects of maltol on D-galactose-induced liver and kidney aging and injury. Phytother Res. 2021;35: 4411–4424. doi:10.1002/ptr.7142

31. Shen L, Fan L, Luo H, Li W, Cao S, Yu S. Cow placenta extract ameliorates d-galactose-induced liver damage by regulating BAX/CASP3 and p53/p21/p16 pathways. J Ethnopharmacol. 2024;323: 117685. doi:10.1016/j.jep.2023.117685

32. Chen Q, Zhao H, Ma N, You X, Yang S, Ma Q, et al. [Decline of secretory function of TM4 Sertoli cells stimulated by D-galactose in mice and its mechanism]. Xi bao yu fen zi mian yi xue za zhi = Chinese J Cell Mol Immunol. 2018;34: 327–333.

